# CRAFT: a bioinformatics software for custom prediction of circular RNA functions

**DOI:** 10.1101/2021.11.17.468947

**Authors:** Anna Dal Molin, Enrico Gaffo, Valeria Difilippo, Alessia Buratin, Caterina Tretti Parenzan, Silvia Bresolin, Stefania Bortoluzzi

**Affiliations:** Department of Molecular Medicine, University of Padova, Padova, Italy; Department of Biology, University of Padova, Padova, Italy; Onco-hematology, stem cell transplant and gene therapy laboratory, IRP-Istituto di Ricerca Pediatrica, Padova, Italy; Pediatric Hematology, Oncology and Stem Cell Transplant Division, Women and Child Health Department, Padua University Hospital

**Author notes:** Corresponding authors Prof. Stefania Bortoluzzi, Dr. Anna Dal Molin.

**Keywords:** CircRNA, bioinformatics, computational pipeline, functional prediction, regulatory network

## Abstract

Circular RNAs (circRNAs), transcripts generated by backsplicing, are particularly stable and pleiotropic molecules, whose dysregulation drives human diseases and cancer by modulating gene expression and signaling pathways. CircRNAs can regulate cellular processes by different mechanisms, including interaction with microRNAs (miRNAs) and RNA-binding proteins (RBP), and encoding specific peptides. The prediction of circRNA functions is instrumental to interpret their impact in diseases, and to prioritize circRNAs for functional investigation. Currently, circRNA functional predictions are provided by web databases that do not allow custom analyses, while self-standing circRNA prediction tools are mostly limited to predict only one type of function, mainly focusing on the miRNA sponge activity of circRNAs. To solve these issues, we developed CRAFT (CircRNA Function prediction Tool), a freely available computational pipeline that predicts circRNA sequence and molecular interactions with miRNAs and RBP, along with their coding potential. Analysis of a set of circRNAs with known functions has been used to appraise CRAFT predictions and to optimize its setting. CRAFT provides a comprehensive graphical visualization of the results, links to several knowledge databases, and extensive functional enrichment analysis. Moreover, it originally combines the predictions for different circRNAs. CRAFT is a useful tool to help the user explore the potential regulatory networks involving the circRNAs of interest and generate hypotheses about the cooperation of circRNAs into the modulation of biological processes.

**Key points:**

- CRAFT is a self standing tool for comprehensive circRNA function prediction.
- CRAFT functions include circRNA sequence reconstruction, microRNA and RNA-binding protein response elements and coding potential prediction.
- Predictions for multiple circRNAs are connected to infer possible cooperation networks and illustrate the potential impact of circRNAs on biological and disease processes.

## Introduction

Circular RNAs (circRNAs) are RNA molecules in which, in a process called backsplicing, a downstream 5’ splice site is covalently linked to an upstream 3’ splice site giving rise to a circle. The backsplice most frequently, although not always, joins canonical exons, with respect to linear transcript annotation [1].

CircRNAs often exhibit cell type- and tissue-specific expression [2,3]. These peculiar RNAs regulate several biological processes with different mechanisms. Moreover, circRNAs are emerging as key oncogenic or tumour suppressor molecules whose expression is dysregulated in disease, cancer and when genomic translocations occur [4–6]. They represent stable diagnostic and prognostic markers [7,8]. Most interesting, the identification of new disease mechanisms involving circRNAs has the potential to indicate novel therapeutic targets. CircRNA functions mostly involve sequence-specific binding with other nucleic acids or proteins, or specific coding potential. One prominent mechanism whereby circRNAs function is by sponging miRNAs, through one or multiple miRNA-response elements (MRE), thus acting as competitive endogenous RNAs (ceRNA) and regulating the expression of miRNA-target genes [9]. Several key oncogenic circRNA-miRNA-gene axes have been described, impacting all cancer hallmarks [10]. With this mechanism, circRNAs also regulate key epigenetic “writers” [11] controlling DNA methylation and histone modifications. CircRNAs can also modulate the activity of RNA-binding proteins (RBP), a large class of molecules involved in most biological processes, since they regulate gene expression at the transcriptional and post-transcriptional levels [12–14]. The protein-RNA interaction is sequence-specific and RBP-response elements (RRE) can be predicted using pattern recognition [15].

Finally, beyond exerting functions typical of long non-coding RNAs, circRNAs can be translated into circRNA-encoded peptides (CEP)[16–18], including circRNA-specific ones generated by translation of open reading frames (ORF) encompassing the backsplice junction, which are not present in linear transcripts, and circRNAs with a rolling ORF, lacking a stop codon a continuing along the circle [19,20] using it as a ‘*Mobius strip*’ [21].

The same circRNA can carry multiple functional sequences, thus playing different functions also with different mechanisms. For instance, for circZNF609 both translation to a peptide and ceRNA activity have been described [17,22], and for circFOXO3 both interaction with proteins and miRNA sponging activity have been reported in literature [23,24].

The function of most of the thousands of circRNAs discovered so far is still undetermined. Therefore, the prediction of circRNA function is instrumental to generate specific hypotheses about their molecular interactions and biological roles to be experimentally investigated, and to interpret the possible impact of circRNA dysregulation in disease.

Several databases have been released with the goal of collecting information regarding known circRNAs. CircBase reports circRNA expression in different biological samples [25], CircR2Disease [26] and Circ2Traits [27] contain, respectively, experimentally validated and potential circRNA association with diseases. CircNet [28] is specific for circRNA-miRNA-mRNA interaction networks, whereas circ2GO [29] merges circRNA expression, gene ontology and miRNA prediction. CircInteractome [30] and CircAtlas [31] collect data about miRNA and RBP binding to circRNAs. RiboCIRC [32] and TransCirc [33] provide information about circRNA coding ability. Importantly, no databases provide predictions of the three above described circRNA functions together. In addition, all of them provide pre-computed data and don’t allow custom predictions. Often, data browsing is difficult and the user has no control about the correspondence between circRNA sequence and functional predictions.

A few tools are available hitherto for circRNA functional prediction. Among them, Cerina R shiny web application [34] and ACT web server [35] predict circRNA functions exclusively based on the ceRNA model. Instead, CircRNAprofiler [36] and CircCode [37] are both dedicated to the coding potential of circRNAs, the second needing ribosome profiling (Ribo-seq) data as input. In summary, all of the available tools allow the prediction of a single function of circRNAs and, to our knowledge, no software is available to comprehensively predict circRNA functions. Moreover, the circRNA sequence is a prerequisite for functional predictions, but most methods to detect circRNA from RNA-seq data return only the backsplice junction coordinates. FcircSEC R package [38] retrieves circRNA sequence using the longer transcript of the gene as reference.

To fill this gap, we developed CRAFT (CircRNA Function prediction Tool), a new computational pipeline to predict circRNA putative molecular interactions with miRNAs and RBP, and circRNA coding potential. Plus, CRAFT allows investigating complex regulatory networks involving circRNAs acting in a concerted way, such as by decoying the same miRNAs or RBP, or miRNAs sharing target genes. This new tool facilitates the interpretation of circRNA biological and pathogenetic roles, and can guide the experimental investigation of circRNA-involving mechanisms.

## Methods

### CircRNA sequence reconstruction

Bedtools v2.28.0[39] was used for file manipulation and sequence extraction, as described in the Results section. The circRNA genomic coordinate file and the build genomic reference file are in BED format. The predicted circRNA sequences are in FASTA format.

To run the predictions circRNA sequences are modified in order to also detect functional elements overlapping the backsplice region. More in detail, a pseudo-circular sequence is created repeating the first 20 nt of the sequence at the end of each circRNA for MRE and RRE prediction, whereas it is repeated twice to detect ORF crossing the backsplice junction. All the prediction coordinates are relative to the circRNA sequence used.

### MiRNA binding site prediction

MiRNA sequences downloaded from miRBase release 22.1 are included in the tool. A filter is then applied to retrieve miRNA of the species selected by the user. MiRanda 3.3a[40] and PITA [41] tools are used to predict miRNA binding sites in circRNA sequences. MiRanda parameters set as default by CRAFT are -sc 80 -en -15; PITA is run with default parameters. AGO2 binding data were downloaded from the doRiNA database [42], merging all 17 files with AGO2 binding information, and converted in hg38 coordinates through UCSC liftOver. Only miRNA binding sites predicted both by miRanda and PITA, and overlapping with AGO2 binding sites, are kept. The AGO2 binding site file is optional. A different file with AGO2 binding data can be provided by the user in BED format.

### RNA binding proteins binding site prediction

RBP binding sites were predicted through beRBP [43] tool, modified to give in output the “voteFrac” score for all predictions, and run in “general mode” searching for all the 143 RBP and 175 Position Weight Matrices of the tool database.

Result tables are given in plain text tabular format, keeping the output similar to that of the tool’s original output, to allow deep inspection to expert users.

### Functional enrichments

The multiMiR 1.8.0 [44] R package is used to perform enrichments of miRNA association with drugs and diseases. Functional enrichments are performed on validated target genes (TG) of miRNAs with predicted binding sites in circRNA sequences. Validated miRNA TG were retrieved from three different databases (miRecords, mirTarBase and Tarbase) using multiMiR, selecting the “strong” validation category of each database. *DOSE 3*.*18*.*2* package [45] is used for over-representation analysis, *clusterProfiler 4*.*0*.*5* package [46] for Gene Ontology analysis and KEGG over-representation test, *ReactomePA 1*.*30*.*0* package [47] for Reactome pathway analysis, *meshes 1*.*12*.*0* package [48] for MeSH enrichment analysis. Functional enrichments are performed also on RBP with predicted binding sites in circRNA sequences using the *UniprotR 2*.*1*.*0* R package [49]; Gene Ontology, KEGG enrichment, Reactome pathway and RBP-disease association analyses are obtained. Protein expression, and functional and miscellaneous information associated with RBP were also retrieved.

### Open reading frames prediction

Putative open reading frames (ORF) are predicted through ORFfinder tool, searching for ORF with the canonical “ATG” start codon, with minimal ORF length of 30 nt, and detected on plus strand. All four kinds of ORFfinder output formats are produced.

### Graphical visualization

The R packages *ggplot2 3*.*3*.*5* [50], *pheatmap 1*.*0*.*12, tidyverse 1*.*3*.*1* [51], *car 3*.*0-11* [52] *and enrichplot 1*.*12*.*2* [53] *are used for result graphical visualization, while tables are printed through the DT 0*.*19* R package.

For the default graphical visualization, CRAFT filters the results with more stringent parameters: miRanda score > 80 and free energy < -15 kcal/mol, PITA ΔG_duplex_ < -15 kcal/mol and ΔG_open_ > -15 kcal/mol are set, beRBP voteFrac ≥ 0.15.

### Assessment and sample analyses

For all the presented analyses, known circRNA backsplice coordinates were retrieved from CircBase. CircRNA binding with miRNAs or RBP, and the coding potential information, were collected from literature (**Table 1**). CircRNA annotation and genomic sequence are based on the Ensembl GRCh38 v.93 human genome. CRAFT was run with default parameters.

**Table 1.** CircRNAs used for the CRAFT assessment analysis, with previously validated functional elements and biological or pathogenetic roles.

CRAFT sensitivity was calculated as true positive predictions (TP, known elements correctly predicted) over the total number of predictions (TP+FN, where FN stands for false negatives). The prediction ratio was calculated as the mean number of predictions at a certain setting over the mean number of predictions obtained at the less stringent condition tested.

## Results

### Overview of the CRAFT workflow

CRAFT is a self standing tool written in Bash and R languages. The Docker [77] containerization makes it usable and portable on all systems.

The CRAFT analysis workflow comprises four main steps (**Figure 1**): (i) circRNA sequence extraction; (ii) circRNA functional predictions, including miRNA and RBP binding sites, and ORF detection; (iii) post-processing of predictions and graphical visualizations of the results for each single circRNA; (iv) integration of predictions for all the circRNA given in input. The first step is performed with the CircExtractor module, which will be described below, in a dedicated paragraph. The functional prediction step relies on the combination of different methods as detailed in **Methods**. The post-processing phase integrates the predictions with knowledge databases for functional enrichment analysis of miRNAs, RBP and the known miRNA target-genes (TG) associated with the circRNAs in the previous step. In addition to the results provided for each circRNA, in the last step, the predictions are combined to identify miRNAs, TG and RBP in common between the different circRNAs.

**Figure 1.**
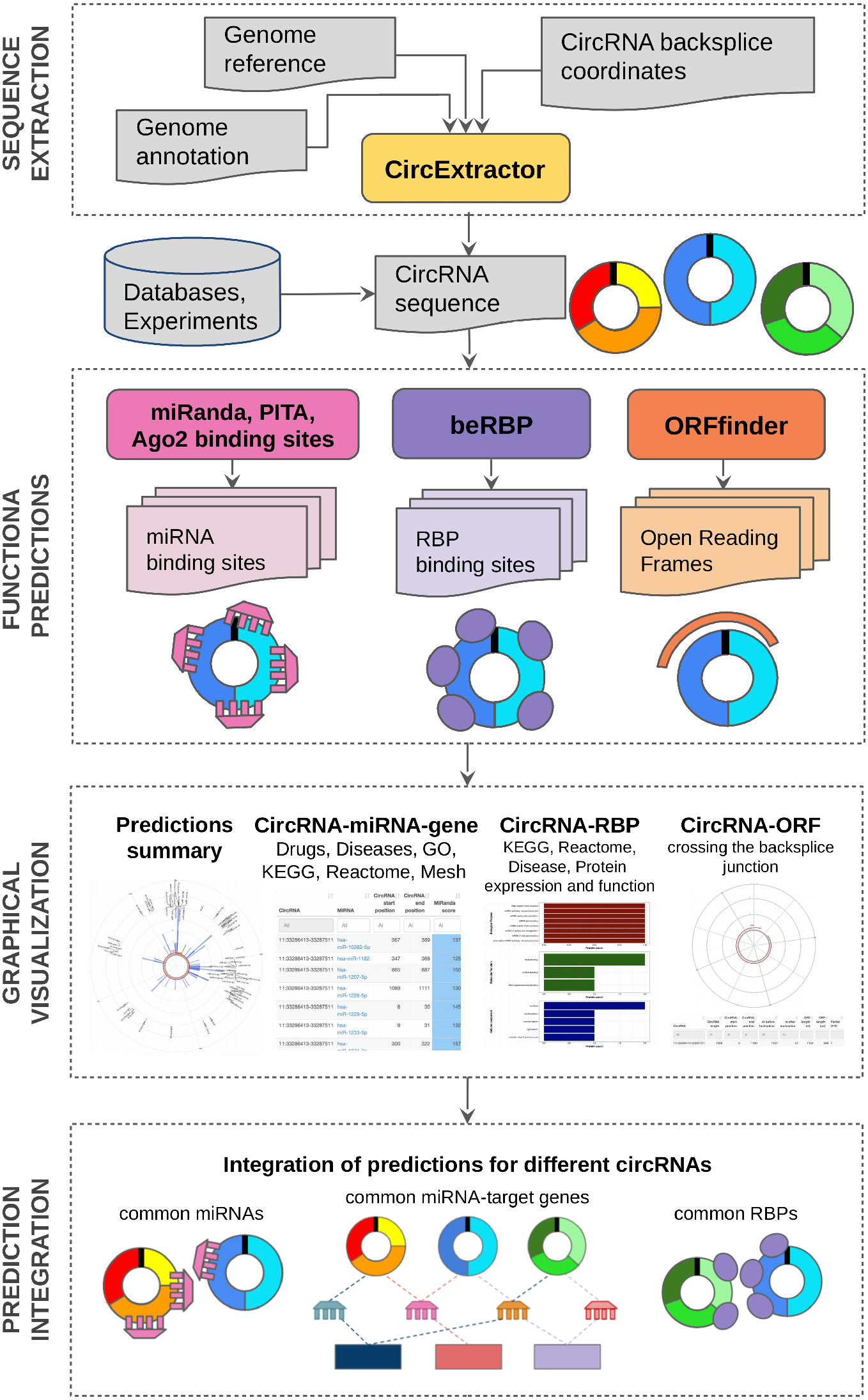
CRAFT workflow, schematizing the pipeline input, the main analysis steps and the produced output.

CRAFT accepts circRNAs of any species: the genome sequence and the gene annotation files must be given in input in FASTA and GTF formats, respectively.

Finally, the user has full control of the parameters used for each tool and can choose to perform either all or only part of the available predictions. Moreover, after the first run, the user can adjust the result visualization by modulating the output filtering parameters without the need of computing the predictions again, saving computational time.

### CRAFT assessment and parameters optimization

We assessed how the parameter tuning and different tool combinations affected the whole number of sites predicted by CRAFT and its capability to detect functional elements, in order to optimize the default setting of the tool. With a literature survey, we collected 20 circRNAs with experimentally determined functions (**Table 1**), including 26 functional elements, that were used as a test set. For each prediction type, we tested whether different strategies and settings can recover the known functional elements in the test set (sensitivity), and how the prediction ratio (see **Methods**) were affected.

The analysis of the known MRE showed that the sensitivity of predictions obtained by miRanda, PITA and intersection thereof is constantly high (≥ 0.9), and the large number of predictions provided by single methods can be controlled by their intersection (**Figure 2A**). The addition of information about experimentally determined AGO2 binding sites, particularly in combination with the intersection of miRanda and PITA predictions, kept in with a good sensitivity (≥ 0.82), while reducing the amount of predictions to only 6%, respective to the total set of predictions obtained from the union of miRanda and PITA. The starting number of predictions to be intersected and combined with AGO2 binding sites is obviously reduced when more stringent thresholds for the four parameters provided by miRanda and PITA are set.

**Figure 2.**
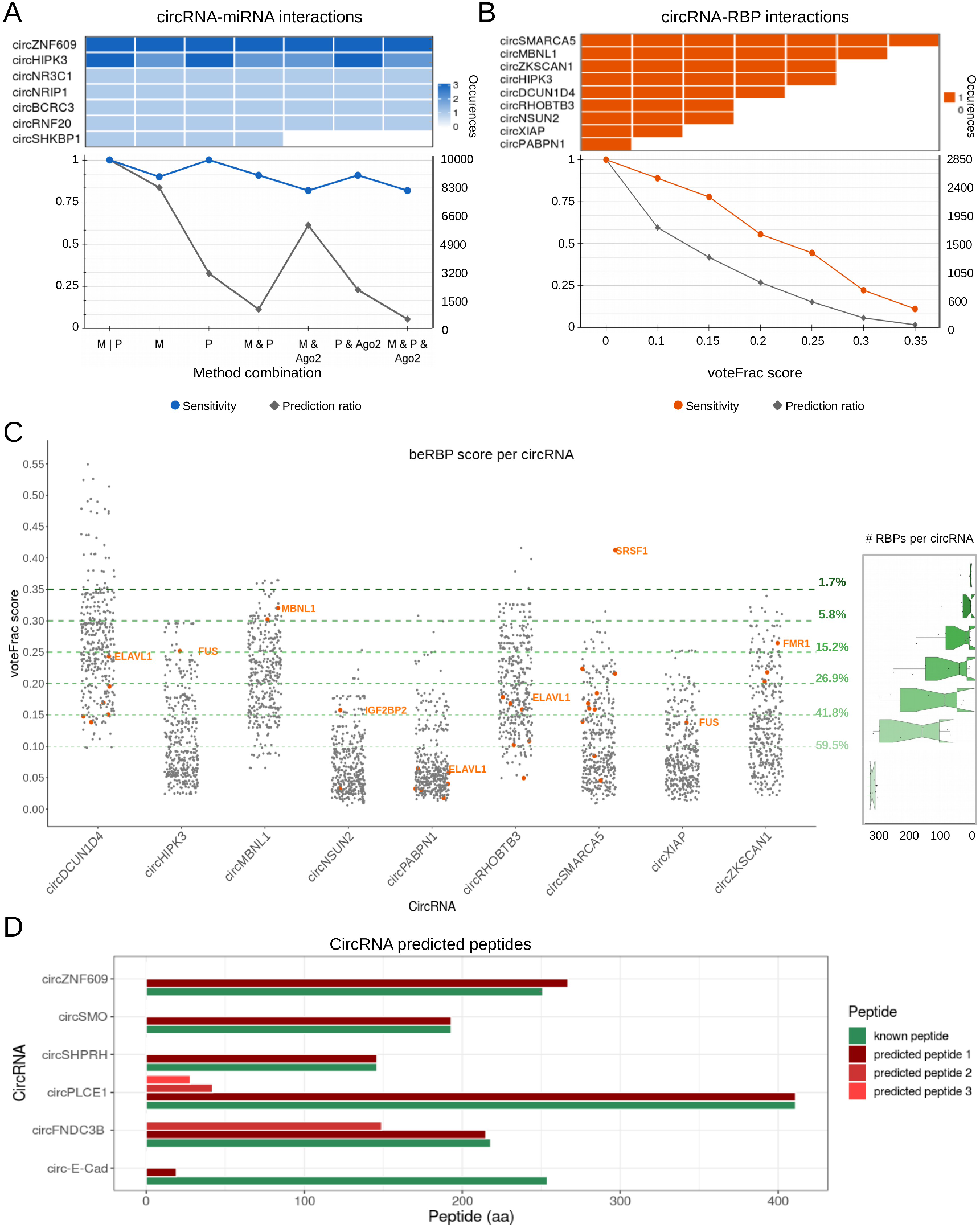
Assessment analysis of circRNAs with known functions. (A) The upper panel shows the number of validated circRNA-miRNA interactions detected by different methods and combinations thereof for each of the considered circRNAs, while the line plot below reports the sensitivity and the prediction ratio (the right axis shows the mean number of predictions for circRNA) in the same conditions (M, miRanda; P, PITA; M | P, union of M and P; M & P, intersection of M and P; the same with AGO2, predicted sites overlapping with experimentally determined AGO2 binding sites); (B) the upper panel shows the number of validated circRNA-RBP interactions detected by different beRBP voteFrac thresholds, while the line plot below reports the sensitivity and the prediction ratio (the right axis shows the mean number of predictions for circRNA) in the same conditions; (C) distribution of RBP binding site prediction scores for each circRNA (known binding sites are shown in orange, the name is shown for the known site with the highest score; the percentage on the right indicates the prediction ratio of the correspondent voteFrac threshold); (D) barplot of the amino acid (aa) length of known (in green) and predicted (in red) peptides for each circRNA.

The scatterplot of prediction scores provided by miRanda (**Supplementary Figure 1**) and PITA (**Supplementary Figure 2**), with indications of the score pairs associated with known MRE, can help the user to understand the effect of stringency on MRE detection. For instance, keeping only the predictions with both scores over the first quartile for both methods, 7/11 known MRE were correctly detected (sensitivity 0.71). Using the median as threshold for the same four scores, the number of predictions decreased (prediction ratio 19%) still identifying 4/11 positives (sensitivity 0.36). To give the user a fairly complete set of predictions, we kept a low stringency on miRanda and PITA parameters in CRAFT default setting, considering that the intersection of miRanda and PITA predictions and the overlapping with AGO2 binding sites controlled for the prediction number, while keeping in with a good sensitivity. Nevertheless, the interactive filtering provided by CRAFT allows the user to customize the number of predictions after inspecting the results.

Regarding RRE, 9/9 of the known TP (sensitivity=1) were detected at the lowest stringency of beRBP predictions score (voteFrac>0), with an average of 316 (range 301-326, median 319) predictions per circRNA. At voteFrac≥0.1, sensitivity was high (0.89), with 8/9 TP detected, and 188 predictions in average per circRNA (range 68-316, median 158) (**Figure 2B**). At increasing stringency (voteFrac≥0.15 and ≥0.2), most of the known functional elements were still detected (sensitivity 0.78 and 0.56, respectively), while considerably reducing the number of predictions to 42% and 27% of the total, respectively (**Figure 2B-C**). At the most stringent score tested, only 1/9 known RRE element (SRSF1 binding circSMARCA5) was detected, being anyhow the unique passing the filter for circSMARCA5, with an average number of 5 predictions per circRNA (range 0-36, median 0). Thus, the best balance between sensitivity and overall number of predictions was observed at a voteFrac≥0.15 (**Figure 2B**), and this optimal threshold was set as default in the CRAFT RBP analysis.

Coding potential analysis predicted an ORF overlapping the backsplice junction for all six tested circRNAs. In 5 out of 6 cases (83.3%), the predicted peptide length exactly corresponded to the validated peptide length (**Figure 2D**). The predicted ORF with different length was that of circ-E-Cad. This was due to the fact that the sequence predicted by CircExtractor based on genome annotation was slightly different from the sequence considered in the previous study. When only the exons from the canonical transcript (Ensembl id: ENST00000261769.10) were used to predict the circRNA sequence, the ORF and peptide were correctly predicted by CRAFT.

### The CircExtractor module

The circRNA nucleotide sequence is necessary to perform functional predictions. Therefore, we included in our pipeline a module that takes as input the circRNA backsplice coordinates, the genome sequence and the gene annotation, and reconstructs the circRNA sequence (**Figure 1**). If circRNA sequences are available to the user, this step can be skipped.

The CRAFT CircExtractor module reconstructs the putative circRNA sequence by intersecting the reference annotation with the backsplice coordinates. Four main cases are considered: (i) if the backsplice joins the ends of two annotated exons, the predicted circRNA sequence contains all and only the exons within the backsplice coordinates (**Figure 3A**); (ii) if one or both backsplice ends map within an exon, only the exon segment within the backsplice coordinates is considered, along with all other entire exons (**Figure 3B**); (iii) if both backsplice ends map in an intronic or intergenic region, the intronic or intergenic part between the backsplice ends is extracted (**Figure 3C**); and (iv), if one backsplice end maps into an exon, whereas the other end maps into an intron or an intergenic region, the sequence will contain all the exons within the backsplice coordinates, plus the intronic/intergenic segment between the backsplice start/end and the first/last exon (**Figure 3D**).

**Figure 3.**
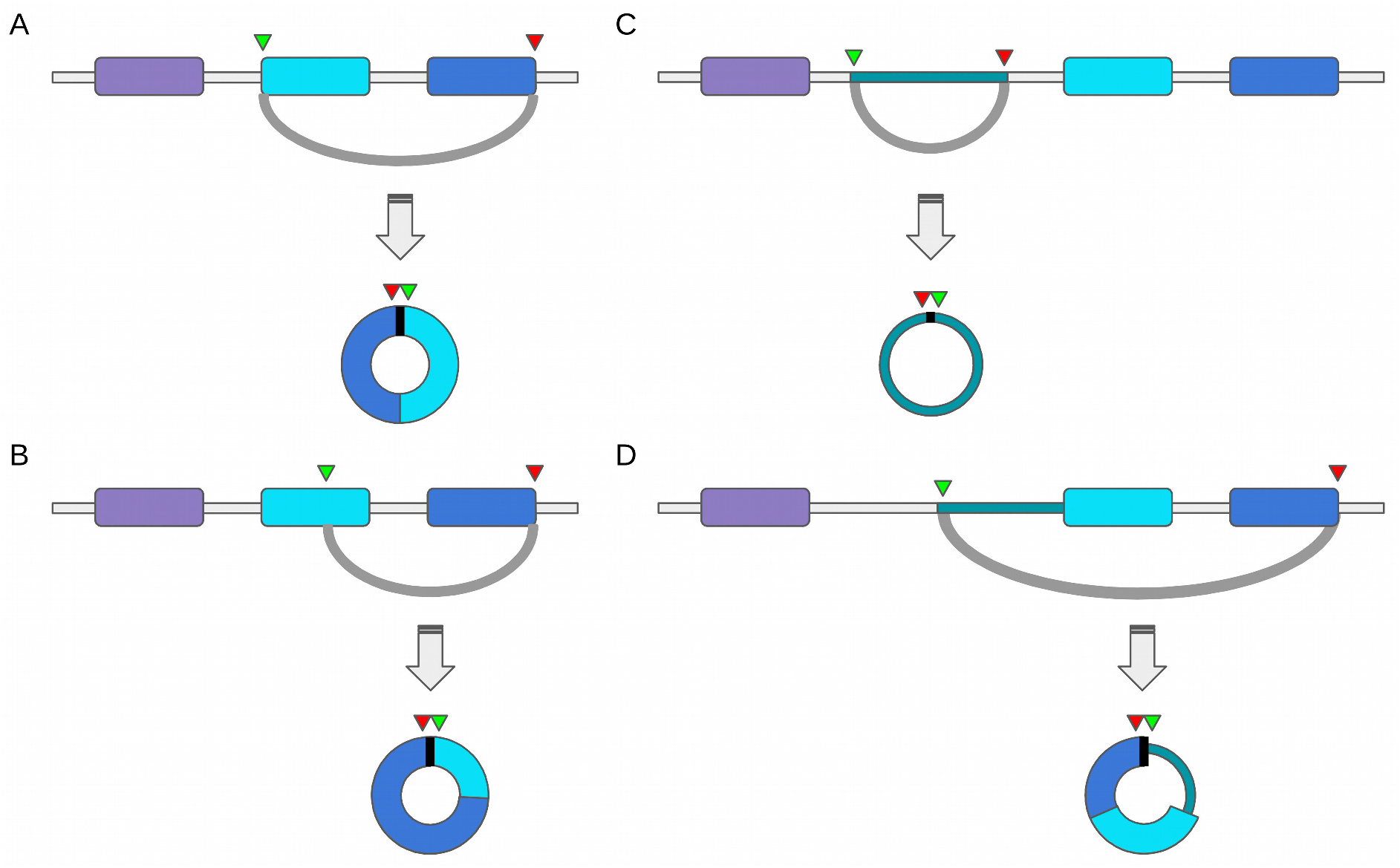
The CircExtractor module of the CRAFT pipeline reconstructs the circRNA sequence using the backsplice coordinates and the genome annotation and reference. (A-D) show the four cases encompassed by the method, according to the position of the backplice ends (green and red triangles represent the backsplice start and end, respectively), in exonic (A-B), intronic (C) and intergenic sequences, and combinations thereof (D).

The reconstructed circRNA sequences are saved in the result directory and automatically passed to the CRAFT functional prediction module.

### CRAFT output

To illustrate CRAFT output, post-processing analyses and data display, we applied CRAFT to three previously characterized circRNAs, circSMARCA5, circHIPK3 and circZNF609, from the above-considered group.

CRAFT outputs multiple HTML pages using interactive tables and figures, describing the predicted miRNA and RBP binding sites, and the putative ORF (**Figure 1**). Prediction result tables link miRNAs, genes and proteins to several databases (miRBase, GeneCards, NCBI Entrez, Ensembl, Uniprot, and NCBI Pubmed). All tables are downloadable in comma-separated value plain text (CSV), Excel, and PDF formats, can be copied to clipboard or printed, allow filtering and ordering, and display coloured bars to highlight important features.

A default set of high-scoring predictions is reported and visualized in the output HTML pages (see **Methods**). The user can tune the filter stringency and re-generate the HTML pages with updated tables and figures with minimal computation time. Two filtering modes are allowed: 1) A “direct mode”, by directly inputting each prediction tool threshold parameter,; 2) A “quantile mode”, to filter a subset of predictions based on a selected quantile (f.i. display and analyze only the top 25% high scoring predictions for each tool).

For each circRNA, a dedicated page is given with detailed predictions, as exemplified by a summary of functional predictions obtained for circZNF609 (**Figures 4-5**). At first, a circular plot shows predicted miRNA and RBP binding positions in the circRNA sequence, and the longest predicted ORF (**Figure 4A**). Next, three sections are dedicated to miRNA, RBP and ORF predictions, respectively.

**Figure 4.**
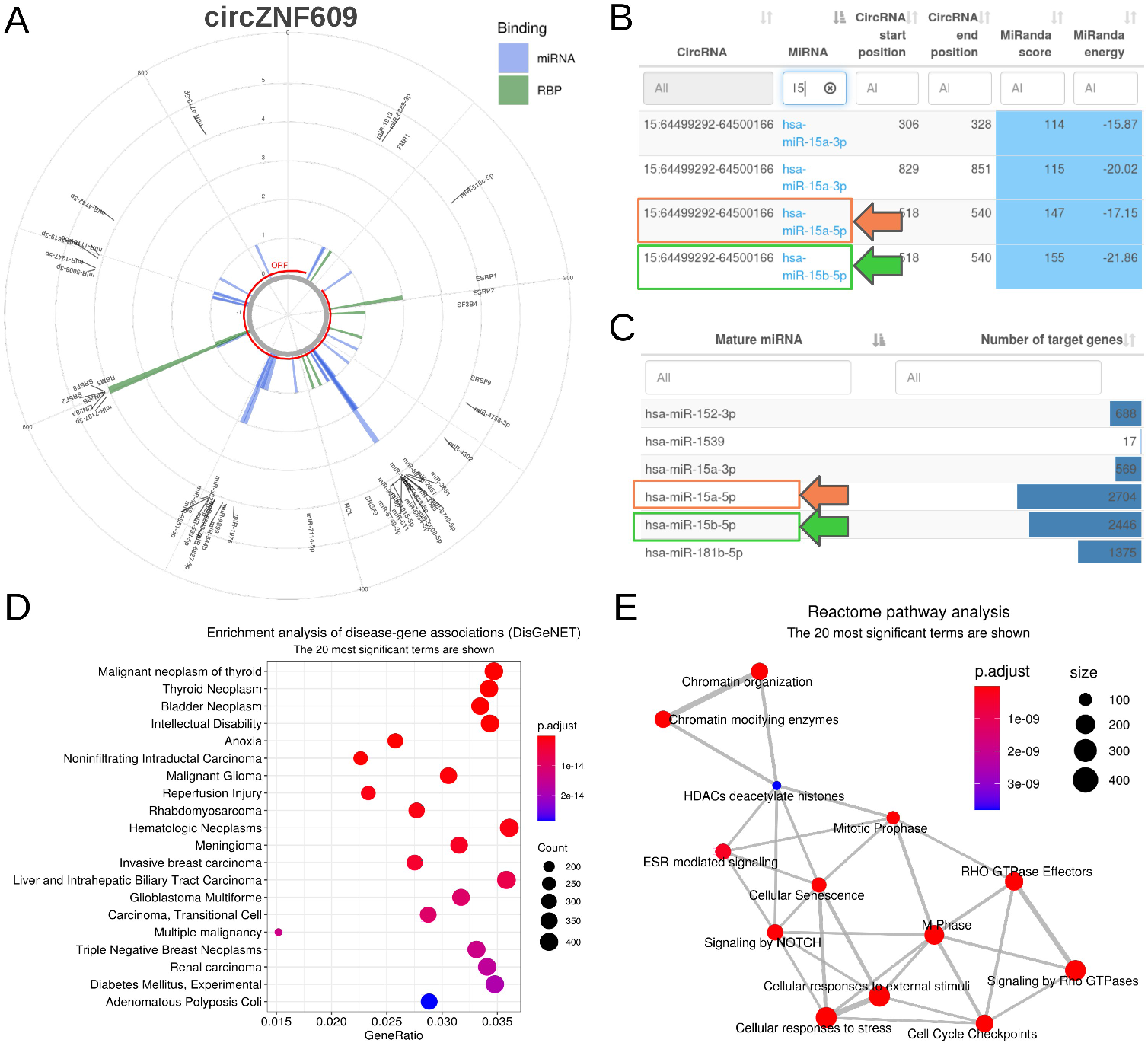
CRAFT miRNA prediction output based on circZNF609 analysis. (A) Summary plot with predicted miRNA and RBP binding sites, and the longest ORF (with a stop codon), along the circRNA sequence; (B) table of CRAFT predicted miRNA binding sites in circZNF609 sequence, with start and end positions, and scores (orange and green boxes highlight predicted miR-15a-5p and miR-15b-5p binding sites, respectively); (C) table summarizing the number of TG per miRNA (box colors as in B); (D) plot of disease-miRNA TG association enrichment analysis; (E) network showing Reactome pathway enrichment analysis results on miRNA TG. In panels B and C only a small part of the corresponding table is shown.

The miRNA section provides, after a circular plot focusing exclusively on miRNA binding sites, a table with the start and end binding positions and the prediction scores (**Figure 4B**). Then, the list of all validated TG is provided for each miRNA, along with different summarization tables, including the number of TG associated with each miRNA (**Figure 4C**), the list of miRNAs associated with each TG, and miRNA-TG interactions associated with drugs or diseases. Functional enrichment results on validated TG are displayed, including disease-associated genes (**Figure 4D**), Gene Ontology, KEGG analysis, Reactome pathways (**Figure 4E**) and MeSH terms. Of note, miR-15a-5p and miR-15b-5p binding sites were detected for circZNF609, in line with literature data [17,61,62].

The RBP section, after a circular plot with RBP binding positions and the corresponding interactive table (**Figure 5A**), displays results of functional enrichments performed on RBP, considering Gene Ontology, KEGG analysis, Reactome pathways (**Figure 5B**) and RBP-disease association (**Figure 5C**). Additional tables provide protein expression, functional and miscellaneous information associated with RBP.

**Figure 5.**
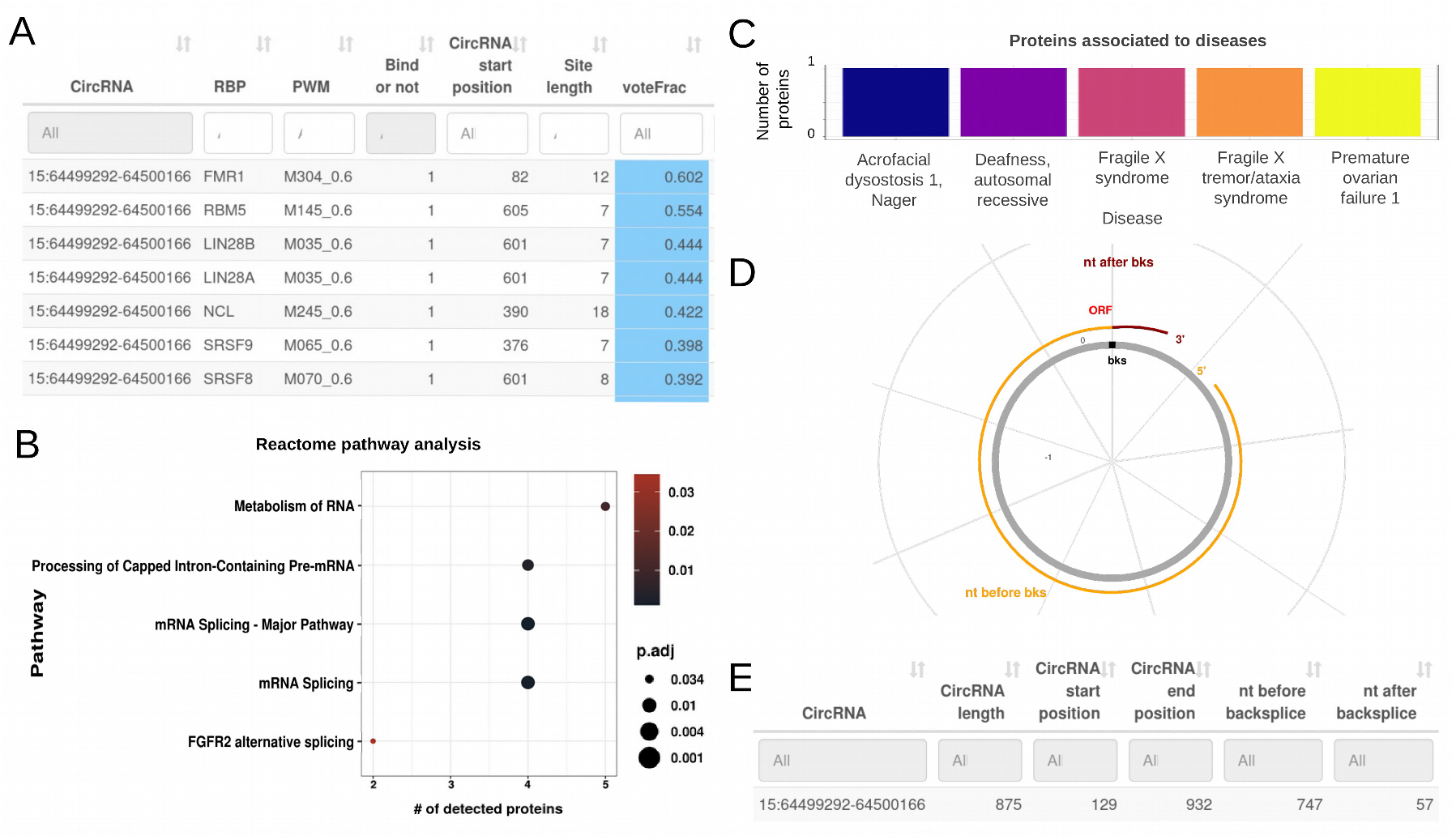
CRAFT RBP and ORF prediction output based on circZNF609 analysis. (A) Table of CRAFT predicted RBP binding sites in circZNF609 sequence, with start position, binding probability and prediction score; (B) plot of Reactome enrichment analyses on RBP potentially interacting with circZNF609, according to CRAFT predictions; (C) barplot of RBP-disease associations; (D) plot of the longest predicted ORF with a stop codon in circZNF609 sequence; (E) table with start and end position, length and frame of predicted ORF. In panels A and E only a small part of the corresponding table is shown.

In the last section, the longest predicted ORF overlapping to the backsplice junction is shown by a circular plot (**Figure 5D**), while a table reports, for all the predicted ORF, the start and end positions, the distance from the backsplice junction, and the predicted coding and peptide sequences in FASTA format (**Figure 5E**).

In principle, different circRNAs can act as sponge for the same miRNA, reinforcing the de-suppression of the miRNA targets. Multiple circRNAs can regulate the same gene also through different miRNAs (**Figure 1**). Thus, if more than one circRNA is given in input, CRAFT integrates altogether the predictions obtained for the different circRNAs. The common predictions are displayed through tables and figures, to provide valuable hints about the possible cooperation of circRNAs to the same regulatory axes. The thirteen miRNAs with binding sites in the top 45% high-scoring predictions for both tools and common to multiple circRNAs, are shown in **Figure 6A**. Moreover, in the CRAFT output, circRNAs are connected to miRNA-validated TG through their predicted interactions with miRNAs. In addition to tables with TG potentially shared by circRNAs, CRAFT provides a network visualization of predicted circRNA-miRNA-TG axes in which multiple circRNAs are connected to the same TG through the same or different miRNAs. For instance, circHIPK3 and circZNF609 can potentially control the expression of a group of eleven genes by sponging miR-1207-5p (circHIPK3) or miR-5581-3p and miR-611 (circZNF609) (**Figure 6B**).

**Figure 6.**
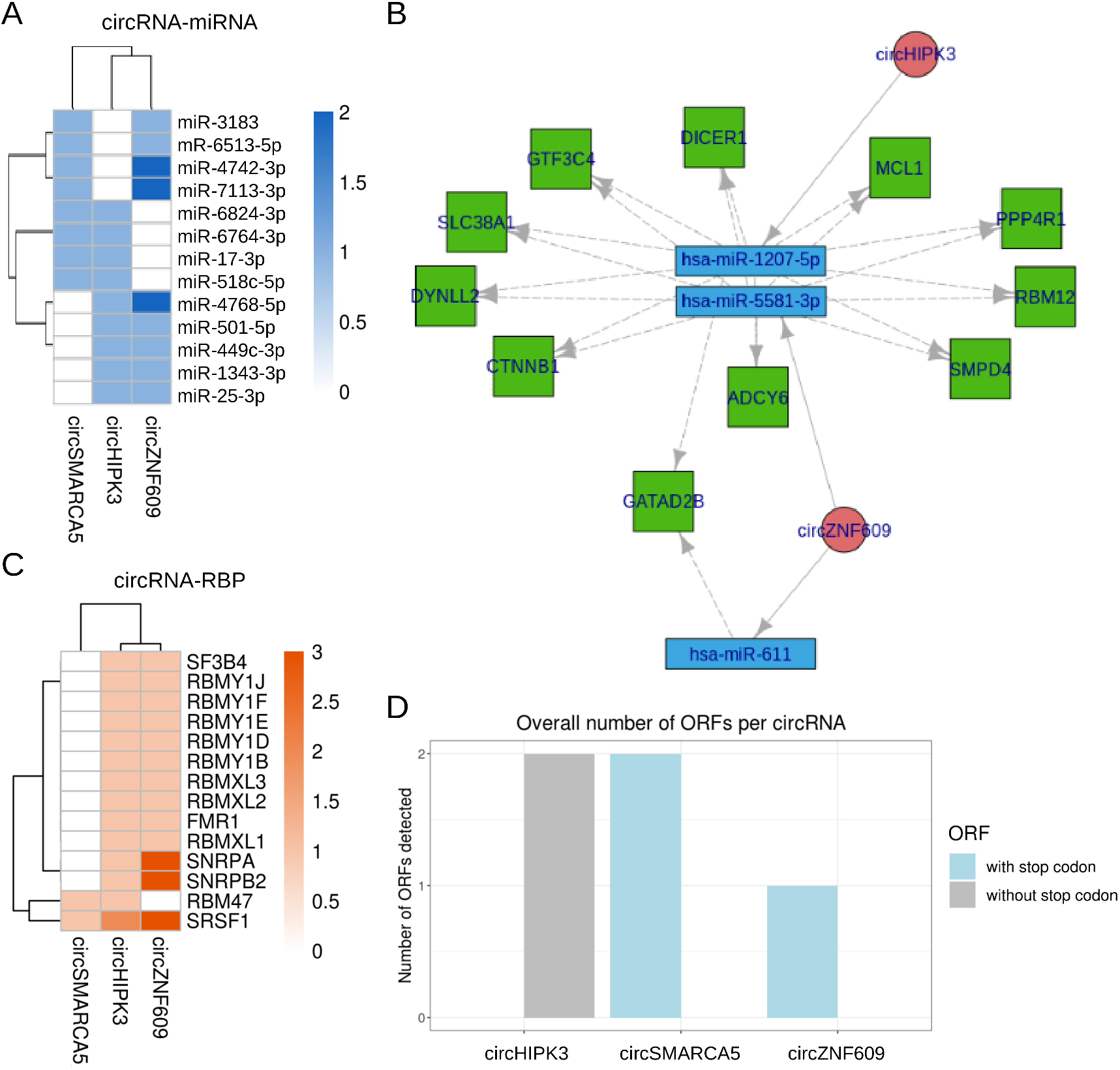
Predictions of different circRNAs are combined to facilitate hypothesis generation. (A) Heatmap of the miRNAs common to at least two circRNAs, considering the top 45% high-scoring miRNA binding site predictions; (B) network showing the eleven common validated TG, targeted by miR-1207-5p, miR-5581-3p and miR-611, respectively associated to circHIPK3 (the first) and circZNF609 (the last two); high scoring predictions was obtained with miRanda score>122 and energy<-24, PITA ΔG_duplex_<-20 and ΔG_open_>-11; (C) heatmap of the RBP common to at least two circRNAs, considering the top 10% high-scoring RBP binding site predictions; (D) summary barplot of the number of ORF predicted for the group of circRNAs considered by CRAFT, binning ORF by the presence or absence of a stop codon in the reading frame. Canberra distance-based clustering has been used for heatmaps in A and C.

Similarly, CRAFT displays RBP associated with RRE predicted in multiple circRNAs. Considering all the predictions obtained with CRAFT default stringency, 48 RBP are commonly associated with all the three circRNAs. Of note, an interaction with the FUS RBP is predicted by CRAFT for circHIPK3, in line with literature data [78], as well as for circSMARCA5. The RBP with binding sites in the top 10% high-scoring predictions and common for multiple of the three circRNAs considered in our sample analysis are shown in **Figure 6C**. All the considered circRNAs have strongly predicted binding sites for SRSF1, including circSMARCA5 whose interaction with SRSF1 has been previously demonstrated [69].

Finally a barplot (**Figure 6D**) and a detailed table present the number of predicted ORF for each circRNA, distinguishing between ORF with and without a stop codon.

## Discussion

Robust evidence on circRNA functions, involvement in biological processes, and oncogenic potential make these molecules extremely attractive for both fundamental and cancer research [79]. CircRNAs can be detected and quantified from RNA-seq data using appropriate software tools, including [80,81]. Next, the challenge faced by studies aiming at defining circRNA roles in disease and cancer is the identification of circRNA-involving mechanisms. CircRNA screening by massive silencing or overexpression studies and experimental investigation can be of great help. Nevertheless, the prediction of circRNA potential interactions and functions is a precondition for the prioritization of circRNAs for experimental functional studies, for the interpretation of experiment results and the design of targeted mechanistic studies. Indeed, even once a circRNA has been demonstrated to significantly impact cell features, such as proliferation, apoptosis, metabolism or differentiation, there is the need to disclose the involved molecules and regulatory axes.

CRAFT has been developed to overcome the limitations of the currently available circRNA function prediction methods. It allows the user to explore putative regulatory networks involving one or more circRNAs of interest, facilitating the interpretation of their biological and pathogenetic role. CRAFT presents four main advantages over the existing tools. First, it provides the putative circRNA sequence. The sequence extraction is based on annotated exons, following the observation that linear splicing patterns of circRNAs mainly join canonical exons [82]. The CircExtractor module in CRAFT merges all the overlapping exons within the circRNA coordinates. This works perfectly well for most cases, as indirectly demonstrated by our test on circRNAs proved to encode for peptides. Of note, CRAFT correctly detected the known ORF of circSHPRH, which is formed by the circularization of four exons of the *SHPRH* gene. In specific instances, the sequence reconstructed by CircExtractor can be different from the sequence produced by the splicing of the exons included in the canonical transcript, as exemplified by circ-E-Cad, whose backsplice ends include a region overlapped by 11 different transcripts that use alternative 5’- and 3’-exon ends in the linear splicing, while the validated circ-E-Cad follows the linear splicing pattern of the canonical longer coding transcript of the gene. In general, the uncertainty associated with circRNA sequence reconstruction based on exon annotations tends to increase with the size of the genomic region within the backsplice ends and, in particular, with the complexity of the alternative linear splicing pattern in the region. In this regard, our recommendation is to use the CRAFT sequence reconstruction module if the circRNA sequences are not known, or for massive analyses of several circRNAs. Nevertheless, in projects focusing on the functional experimental investigation of one or a few specific circRNAs, we advise using experimentally determined circRNA sequences, to provide a firmer ground to build predictions on.

The second main CRAFT asset is that it allows prediction of the three most important circRNAs functions, whose biological relevance is supported by robust literature [9– 14,16,18–20]. The most investigated function of circRNAs is the ability to bind miRNAs. One limitation of miRNA binding site prediction algorithms is the low specificity of the predictions. To face this problem, we used a dual strategy: (i) the combination of two algorithms, miRanda[83] and PITA[84], and (ii) the integration of predictions with experimentally determined AGO2 binding sites. The assessment of the CRAFT predictions on a sizable set of circRNAs with known functions showed that this approach can control the number of predictions, while holding good sensitivity.

Also the interaction of circRNAs with RBP can be associated with important functions [23]. In principle, circRNAs acting as decoys can counteract RBP functions, whereas in other cases circRNAs can serve as scaffolds for the assembly of molecular complexes. CRAFT allows inspecting the possible interactions between circRNAs and RBP, and also the identification of the RBP possibly bound by multiple circRNAs. Of note, beyond the beRBP database of position weight matrices embedded in CRAFT, custom matrices can be provided by the user, extending the analysis.

CRAFT predicts the ORF in the circularized sequence, allowing the user to obtain the peptides potentially encoded by the circRNA and not by the linear transcript. To what extent circRNAs are translated it is still a matter of debate [85], but circRNA-encoded peptides have been identified in different studies and, in several cases, were also proven to be biologically active [86–88]. CRAFT aims at the prediction of circRNA-specific peptides, thus it focuses on those ORF overlapping the backplice site, which are different from the ORF observed in linear transcripts. To undergo translation without a 5’ cap, circRNAs can require specific sequence elements (internal ribosome entry sites (IRES) [89] and N^6^-methyladenosine modification [18,90]) whose prediction can be envisaged as an as interesting CRAFT future development.

The third CRAFT advantage for the user is that it provides a rich output with graphical visualizations, leveraging functional enrichments, and results linking to several knowledge databases. The output can be explored, filtered and re-analyzed as long as the user prefers to generate hypotheses. For instance, the connection of circRNAs with miRNAs, validated miRNA-target genes, and specific pathways, functions and diseases can be of great help to identify regulatory networks to be further scrutinized. This can be pursued with experimental investigation, as well as integrating CRAFT results with circRNA and gene expression data. For instance, positive correlations between a circRNA and the validated target genes of miRNAs potentially sponged by the circRNA can be indicative of functional regulatory axes [4,5].

Finally, we believe that the combination of predictions for different circRNAs provided by CRAFT is a highly useful and original feature. In research projects, when circRNAs dysregulated in cancer or disease tissue compared to a normal counterpart are identified, subsequent experimental investigation often focuses only on the elucidation of the functions of one specific circRNA. Instead, cooperative effects of multiple dysregulated circRNAs in pathogenetic mechanisms can be hypothesised. Thus, the potential interaction of different circRNAs with the same RBP can be inspected in CRAFT. Plus, CRAFT allows the exploration of regulatory networks in which multiple circRNAs can act as sponge for the same miRNA, reinforcing the de-suppression of the miRNA-target genes, or sponge different miRNAs with a common target gene, regulating its expression.

CRAFT is also a highly portable tool, thanks to the containerization of the software, easy to use, and it requires minimal input data. In this study, we appraised CRAFT predictions by using, as benchmarking, a set of circRNAs with known functions, including the three types of functional elements predicted by CRAFT. CRAFT sensitivity was assessed in relation to the total number of predictions to be handled by the user at different settings. Our sensitivity estimates were highly conservative, based on a pejorative scenario in which all the functional elements are known and thus all the predictions not corresponding with known functional elements are counted as false positives, even if some of them can be functional still not known. The conducted tests informed the effects of different parameters, allowing us to optimise the CRAFT strategy and default settings and providing intelligence about the analysis customization and results filtering for CRAFT users.

In conclusion, we are handing over the researchers in the circRNA field with a new software tool facilitating the exploration of circRNA functions, and indicating potential regulatory axes involving one or more circRNAs of interest. The generation of new hypotheses about the potential impact of circRNAs in biological processes, pathways and diseases can help circRNA prioritization for further study and the interpretation of their biological and pathogenetic role.

## Data Availability

CRAFT tool is available on GitHub at https://github.com/annadalmolin/CRAFT and on DockerHub at https://hub.docker.com/repository/docker/annadalmolin/craft.

## Acknowledgements

This work has been supported by AIRC, Milano, Italy (Investigator Grant – IG 2017 #20052 to SB), Italian Ministry of Education, Universities and Research (PRIN 2017 #2017PPS2X4_003 to SB), Fondazione Umberto Veronesi, Milano, Italy (to EG).

## Author Contributions

Conceptualisation, AD and SBo; Data curation, AD; Formal Analysis, AD and SBo; Funding Acquisition, EG, SBr and SBo; Methodology, AD, EG; Project Administration, SBo; Resources, SBo; Software, AD, VD; Supervision, EG and SBo; Visualization, AD and SBo; Writing – Original Draft, AD, EG and SBo; Writing – Review & Editing, AD, EG, AB, CTP, SBr and SBo. All authors read and approved the final manuscript.

## Short description of the Authors

Anna Dal Molin is post-doc at the Computational Genomics Laboratory at the Department of Molecular Medicine, University of Padova. Her research interests include bioinformatics and transcriptomics, circular RNAs in leukemias, and development of computational methods for circular RNA function prediction.

Enrico Gaffo is post-doc at the Computational Genomics Laboratory at the Department of Molecular Medicine, University of Padova. His research interests include circular RNA, microRNA, advanced methods for RNA-seq data analysis, and bioinformatics applied to cancer research.

Valeria Difilippo graduated in Industrial Biotechnologies at the University of Padova working on circRNA prediction for her master thesis.

Alessia Buratin is PhD student in Biosciences (curriculum Genetics, Genomics and Bioinformatics) of the University of Padova. Her main interests are biostatistics and bioinformatics, transcriptomics of hematologic malignancies, and circular RNA biogenesis.

Caterina Tretti Parenzan is a PhD student in Pediatric Oncoematology of the University of Padua. Her research is mostly focused on the study of the oncogenic molecular mechanisms involving circRNAs in pediatric acute lymphoblastic leukemia.

Silvia Bresolin is assistant professor of Molecular Biology at the Department of Women and Child Health Department of the University of Padova. Her field of interest is focused on pediatric leukemia transcriptomics and genomics alterations, as well as functional genomics and development of novel disease models.

Stefania Bortoluzzi is associate professor of Applied Biology at the Department of Molecular Medicine of the University of Padova, where she leads the Computational Genomics Laboratory. Her research interests include cancer genomics and transcriptomics, bioinformatics, systems biology, non-coding RNAs, circular RNAs, exosomal RNAs, and hematologic malignancies.

## Supplementary Materials

**Supplementary Figure 1. Prediction scores provided by miRanda**. Scatterplots and boxplots of prediction scores provided by miRanda (*miRanda score* and *miRanda energy*). Best predictions are in the top right corner (higher *score* and lower *energy*). Known binding sites are shown in blue.

**Supplementary Figure 2. Prediction scores provided by PITA**. Scatterplots and boxplots of prediction scores provided by PITA (*PITA ΔG*_*duplex*_ and *PITA ΔG*_*open*_). Best predictions are in the top right corner (lower *ΔG*_*duplex*_ and higher *ΔG*_*open*_). Known binding sites are shown in blue.

