## Supplementary Materials for "CRAFT: a bioinformatics software for custom prediction of circular RNA functions"

**Supplementary Figures**

Supplementary Figure 1………………………………………………………………………………………..2

Supplementary Figure 2………………………………………………………………………………………..3

**Supplementary Figure 1. Prediction scores provided by miRanda**

Scatterplots and boxplots of prediction scores provided by miRanda (*miRanda score* and *miRanda energy*). Best predictions are in the top right corner (higher *score* and lower *energy*). Known binding sites are shown in blue.

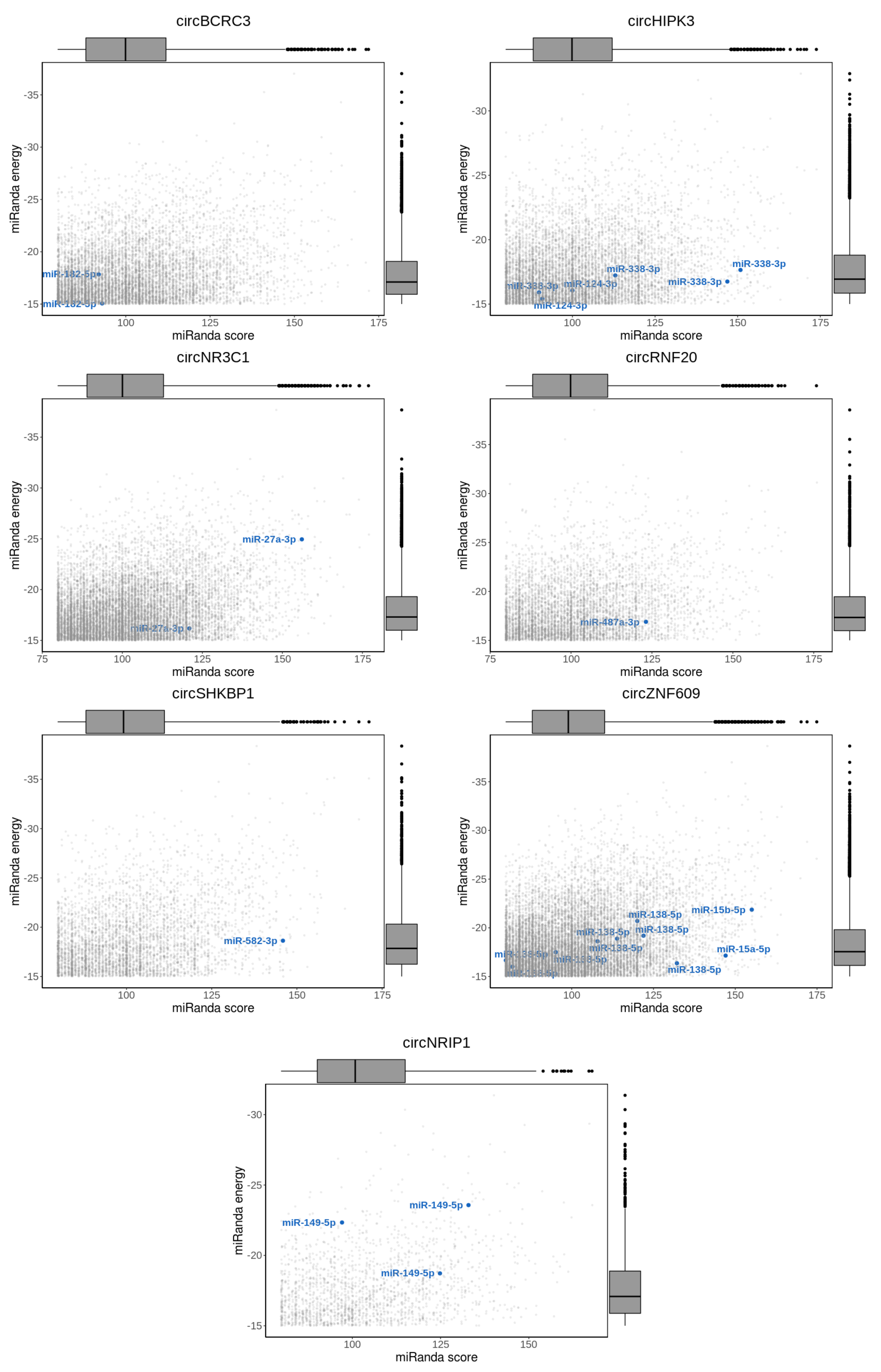

**Supplementary Figure 2. Prediction scores provided by PITA**

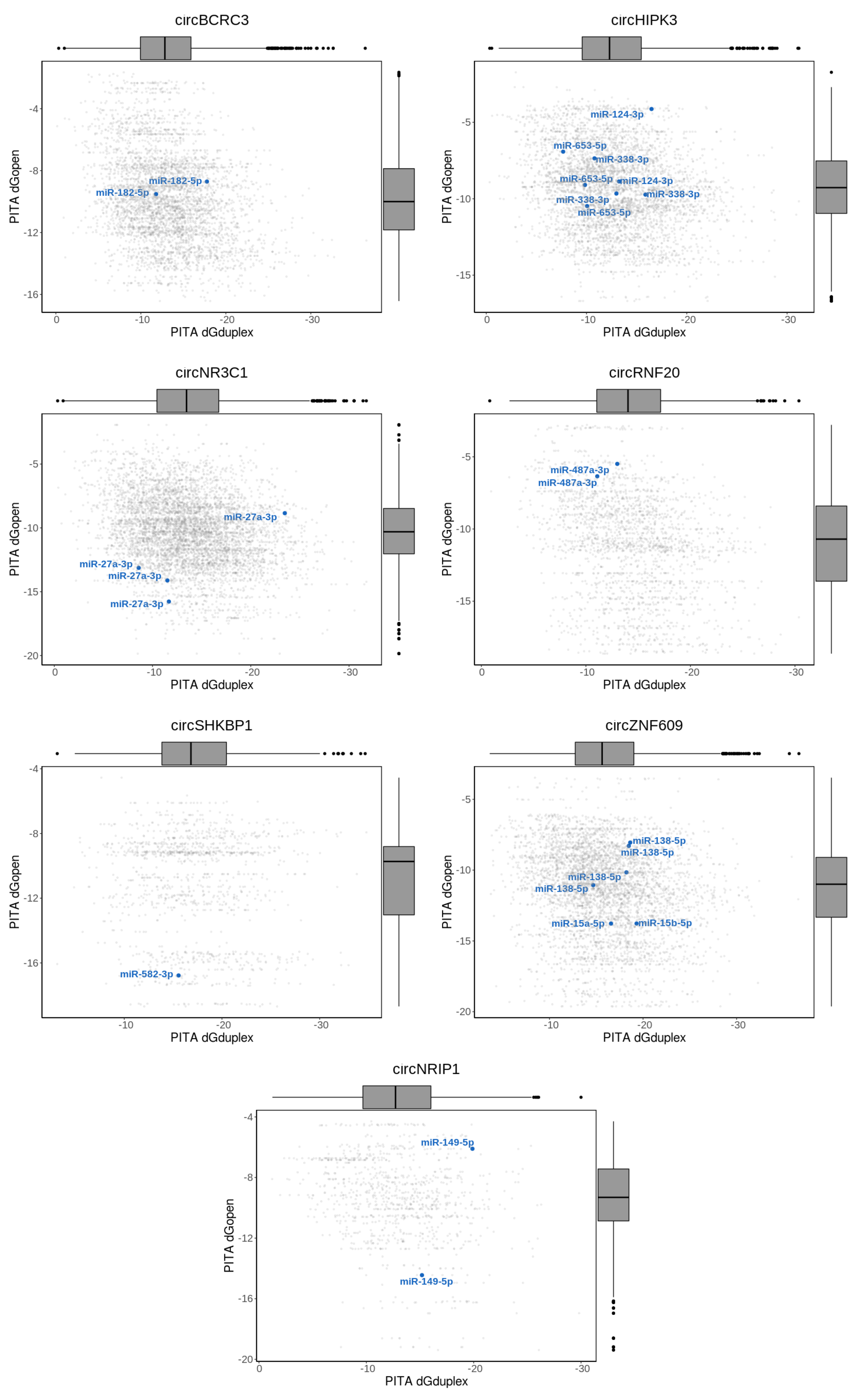

### Scatterplots and boxplots of prediction scores provided by PITA (*PITA ΔGduplex* and *PITA ΔGopen*). Best predictions are in the top right corner (lower *ΔGduplex* and higher *ΔGopen*). Known binding sites are shown in blue.
